# Tissue-like compression stiffening in biopolymer networks induced by aggregated and irregularly shaped inclusions

**DOI:** 10.1101/2025.06.02.657447

**Authors:** Xuechen Shi, Jordan L. Shivers, Fred C. MacKintosh, Paul A. Janmey

## Abstract

Biological tissues experience mechanical compression under various physiological and pathological conditions and often exhibit compression stiffening, in which their stiffness increases during compression, a phenomenon that plays a crucial role in regulating cell behavior and maintaining mechanical homeostasis. However, most isolated biopolymer networks, such as fibrin and collagen hydrogels that form the extracellular matrix and actin network that forms the internal cytoskeleton, undergo compression softening, raising questions about how tissues achieve compression stiffening despite the softening properties of their extracellular and intracellular matrix components. Previous studies have shown that spherical inclusions at large volume fractions can induce compression stiffening in biopolymer networks, but they do not account for the effects of aggregation and irregular morphologies of cellular assemblies or other components in tissues. Here, we demonstrate a novel mode of compression stiffening induced by aggregated or irregularly shaped inclusions that occurs at significantly lower volume fractions. Using carbonyl iron particles and coffee ground particles, we find that the morphological diversity of inclusions enables tissue-like compression stiffening at a low volume fraction of inclusions. Through a set of experiments and computational analyses, we demonstrate that these particles can percolate at low volume fractions. We further show that the percolation of stiff inclusions creates a stress-supporting network and enables tension-dominated stress propagation in fibrin fibers, both of which drive macroscopic stiffening during compression. These findings provide insights into the regulation of tissue stiffness and have implications for designing biomaterials with physiologically relevant mechanical properties for biomedical applications.

**Significance Statement:** Biological tissues experience a variety of mechanical forces. Many tissues, such as brain, liver, fat, and blood clots, become stiffer under physiological compressive loads, a property known as compression stiffening. In contrast, most biopolymer networks, which are the primary structural components for tissues, soften under compression. Here, we show that incorporating a small amount of aggregated or irregularly shaped particles into biopolymer gels induces robust compression stiffening. These inclusions percolate through the gel and rearrange non-affinely under compression, stretching surrounding fibers and contributing to mechanical reinforcement. Together, these effects reproduce tissue-like compression stiffening. Our findings not only provide new physical models for understanding tissue mechanics but also offer insights for designing biomaterials to achieve physiologically relevant mechanical responses.

## Introduction

Many biological tissues, such as the brain (1, 2), blood clots (3, 4), liver (5), and fat (6, 7), experience mechanical compressions in physiological or pathological conditions. These tissues typically exhibit increased stiffness under compression, a phenomenon known as *compression stiffening* (8, 9). For instance, while healthy brain tissue is very soft (shear modulus ∼ kPa), conditions such as tumors and traumatic brain injuries can compress the brain and stiffen the tissue (1, 10-13), which in turn alters cell functions (1). Similarly, gluteus adipose tissues in the buttock region undergo significant compression when individuals sit (14, 15), resulting in tissue stiffening (16). Tissue stiffness is a key biological characteristic that not only provides structural support for physical activities and resistance against gravity, but also regulates cell behaviors through mechanotransduction pathways (17-20). Since tissue stiffness can be dynamically altered by compression from local solid stress or fluid pressure, understanding the mechanisms of compression-stiffening is essential for comprehending tissue mechanics and developing strategies for material design in tissue engineering. However, the primary structural components of tissues, such as collagen and fibrin networks alone as well as cytoskeletal networks formed by actin filaments or microtubules, do not exhibit this compression-stiffening response and tend to soften under compression (21, 22).

To address this difference between fibrous polymer networks and biological tissues, studies have identified two mechanisms involving the inclusion of either soft but volume-conserving or stiff, spherical cell-mimicking particles (9, 23, 24). When a biopolymer hydrogel composite is compressed, such inclusions suppress fiber buckling and promote local fiber stretching, resulting in macroscopic compression stiffening. For soft inclusions, their incompressibility under compression generates localized outward forces that place nearby fibers under tension. For stiff inclusions, their resistance to deformation causes them to rearrange nonaffinely within the matrix, similarly stretching adjacent fibers.

While these models explain modes of compression stiffening, they have primarily focused on monodisperse, uniformly distributed spherical inclusions. However, in biological tissues, cellular components often exhibit irregular shapes due to cell spreading and migration (25), and form aggregates with diverse morphologies. For example, in blood clots, platelets adhere and aggregate to seal wounds and stop bleeding. Upon activation, platelets extend pseudopods, interacting with each other and with fibrin fibers to create a dense three-dimensional network that enhances clot stability (26). Red blood cells can also become wrapped by fibrin fibers, forming tightly packed aggregates that contribute to the mechanical properties of clots (27). Moreover, metastatic tumor cells aggregate and proliferate at metastatic sites, forming clusters of various sizes and irregular shapes due to the absence of E-cadherins (28, 29). These microtumors or micrometastases interact with the extracellular matrix (ECM) in multiple ways that influence tissue mechanical properties (30, 31). Other complex examples include cells cultured in fibrous networks in vitro, which induce compression stiffening at lower volume fractions than monodisperse spheres (9). Beyond cellular components, non-cellular structures with highly irregular geometries in tissues, such as cholesterol-containing crystals in steatotic liver, have been shown to enhance compression stiffening under physiological loads (32). Similarly, studies using irregularly shaped starch particles have shown tissue-like mechanical responses at lower volume fractions than those reported for spherical inclusions (33). These observations indicate that biological tissue stiffening may arise not only from the presence of inclusions but also from their shape, interactions, and aggregation behaviors.

In this study, we investigate the mechanism of compression stiffening in biopolymer networks induced by cell-mimicking aggregated particles and irregularly shaped particles. We show that the inclusion of these particles induces robust compression stiffening in fibrin and collagen gels, even at low volume fractions, a behavior that is not predicted by previously reported models. Our results emphasize the importance of diverse morphologies of these inclusions, rather than polydisperse sizes alone, in achieving tissue-like mechanical properties in the biopolymer hydrogel composites. We further demonstrate that these inclusions form a percolated network at low volume fraction, facilitating tension-dominated stress propagation through the fibrous network as well as providing a stress-supporting structure that enhances macroscopic stiffening under compression. Our findings suggest that non-ECM components may play a significant role in determining tissue mechanical properties, even when the ECM properties are essential for tissue stiffness. Additionally, these insights provide guidance for designing biomaterials with physiologically relevant mechanical properties for applications in tissue engineering and regenerative medicine.

## Results

We first synthesize fibrin hydrogels with 5 mg/ml fibrinogen (∼0.8% volume fraction) embedded with carbonyl iron particles (CIPs). CIPs have a native diameter of 2-8 μm and exhibit slight aggregation when dissolved in aqueous solutions (Fig. S1). During fibrin gel formation, these particles aggregated, forming structures of varying sizes within the fibrin matrix (Fig. 1A, Fig. S2) that are significantly larger than those formed in solutions. Similar CIP aggregation has been observed in other gels (34, 35). The mechanism behind this aggregation remains unclear but might be related to the adhesion of the CIP to growing fibrin polymers during gelation. CIPs remain stable within the fibrin hydrogels and do not oxidize for at least two months (Fig. S3).

**Figure 1.**
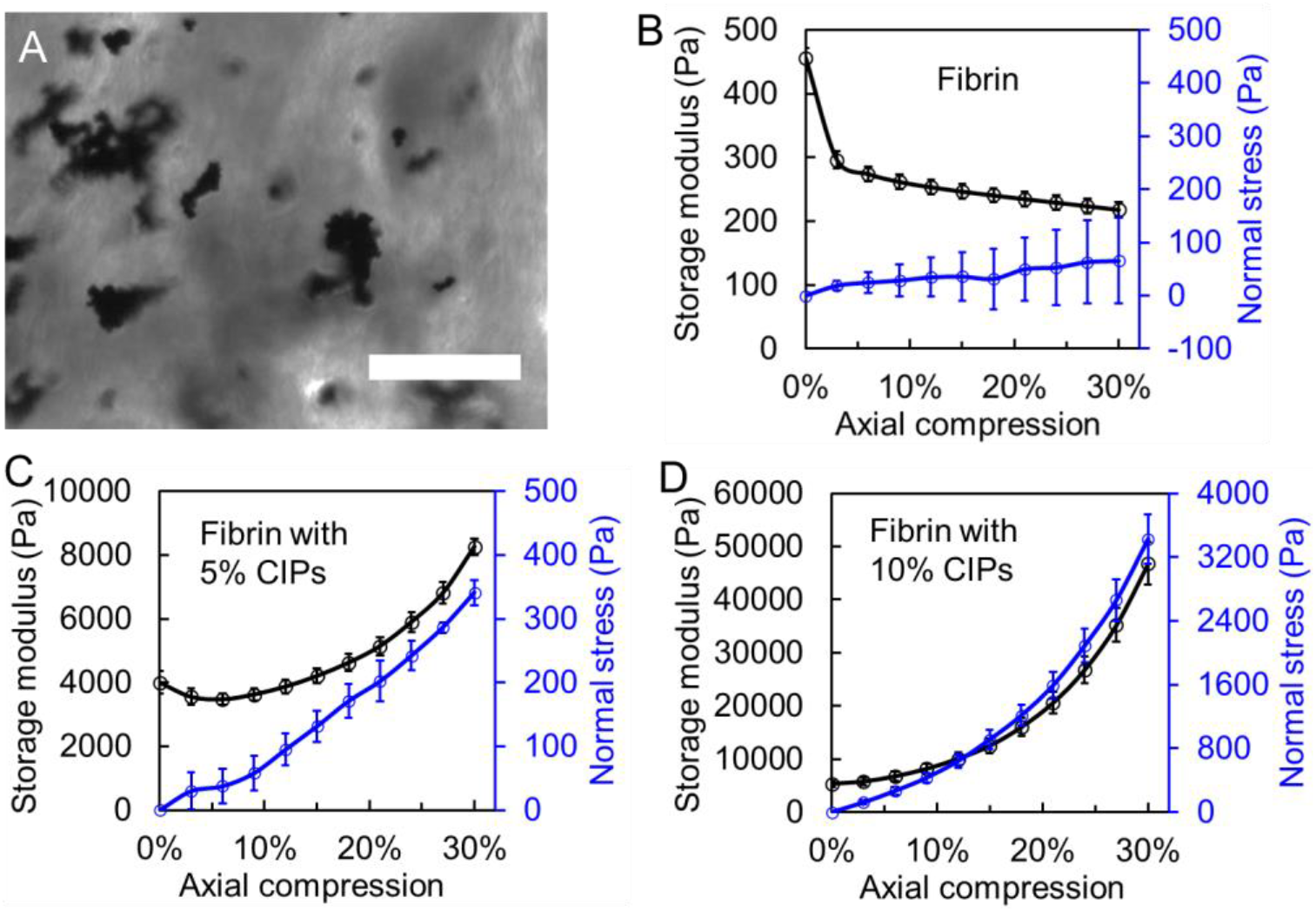
Compression stiffening of fibrin gels with CIP inclusions. **A.** Bright-field image of CIP aggregates in a three-dimensional fibrin network. The volume fraction of CIPs is 1%, which ensures adequate light penetration for imaging. CIPs spontaneously aggregate during the fibrin gelation process. Scale bar = 50 μm. **B-D**. Shear storage modulus and axial stress as a function of axial compressive strain for fibrin gels with different CIP volume fractions: 0% (B), 5% (C), and 10% (D). n≥3 for all curves. Error bar denotes SEM.

The stiffness of the fibrin hydrogels was determined by dynamic shear rheometry, utilizing a 1% strain at 1 Hz. Consistent with previous studies using stiff particles such as dextran beads, the inclusion of CIPs increases the stiffness of the fibrin gels, yet the increase caused by CIPs is notably more prominent, e.g., from 0.5 kPa to 5.5 kPa with 10% volume fraction CIPs. We attribute this enhancement to the matrix-densification-around-inclusions mechanism, as reported previously (36) (Supplementary Note I and Fig. S4), where a similar 10 fold-increase in stiffness has been observed in various types of hydrogels with inclusions at 10% volume fraction. In addition, although not directly related to our system, a recent study demonstrated that embedding prestressed structures within soft hydrogels can result in dramatic increases in stiffness and resilience (37).

Next, we subjected the fibrin gels to stepwise uniaxial compression (setup shown in Fig. S5), increasing by 3% increments up to 30% total compression, and performed small-strain oscillatory shear measurements at constant levels of compression. Incorporation of CIP aggregates, even at low volume fractions (e.g., 5%), transforms the mechanical behavior of the fibrin gel from compression softening to compression stiffening (Fig. 1B-D). Specifically, we observe a significant increase in shear modulus after 30% compression, with a 2-fold or 4 kPa increase for 5% v/v CIPs and a 9-fold or 40 kPa increase for 10% v/v CIPs. For gels with 5% v/v CIP inclusion, an initial softening occurs during compression up to 12%, followed by a transition to stiffening (Fig. 1C). This behavior aligns with the fiber-stretching-around-stiff-inclusions mechanism described in a previous study (23) at small compression levels. To test the generality of our findings, we demonstrate that 10% carbonyl iron particles also transform a collagen network from compression softening to compression stiffening (Fig. S6).

While most fibrous biopolymer networks exhibit shear strain stiffening (38-40), biological tissues are typically shear softening (5, 9, 41, 42). This property is replicated by the fibrin gel embedded with CIPs (Fig. S7): without CIPs, pure fibrin stiffens 5.4 times at 100% shear strain, whereas fibrin with CIPs softens by 65% at 30% shear strain. These shear stiffening/softening properties are maintained even when the gels are subjected to 30% axial compression. Together, these results demonstrate that the inclusion of CIPs in the fibrin network, even at very volume fractions, induces tissue-like mechanical responses, significantly altering the mechanical behaviors of the network under both shear and compressive strains.

We next explore the possible mechanisms of the tissue-like compression stiffening in fibrin gels. The established mechanism of fibers stretching around stiff inclusions (23, 24) requires a volume fraction exceeding 30% to achieve significant stiffening, whereas we observe that a 5% volume fraction of CIP aggregates induces stiffening. Possible differences between our results and previous studies include different sizes of particles, polydispersity, and non-spherical shapes. We test the size effect of rigid particles using monodisperse polystyrene (PS) beads with a diameter of 3 μm or 20 µm within the fibrin gel. Like CIPs, 10% PS beads increase the initial shear modulus of fibrin before compression, but unlike CIP aggregates, a 10% volume fraction of PS beads, whether 3 μm or 20 μm, does not alter the compression softening property of pure fibrin (Fig. 2A, Fig. S8), suggesting that the observed compression stiffening is not attributable to the smaller size of the CIPs. Additionally, it is possible that particle aggregation may induce pre-stress in the fibrin fibers. The shear strain softening response (Fig. S7) also supports this pre-stress condition, as a previous study showed that pre-stressing the fibrin network abolishes shear strain stiffening or induces shear strain softening (9). However, that study also demonstrated that pre-stress alone does not induce compression stiffening. Therefore, we conclude that prestress is not the underlying mechanism for the observed compression stiffening.

**Figure 2.**
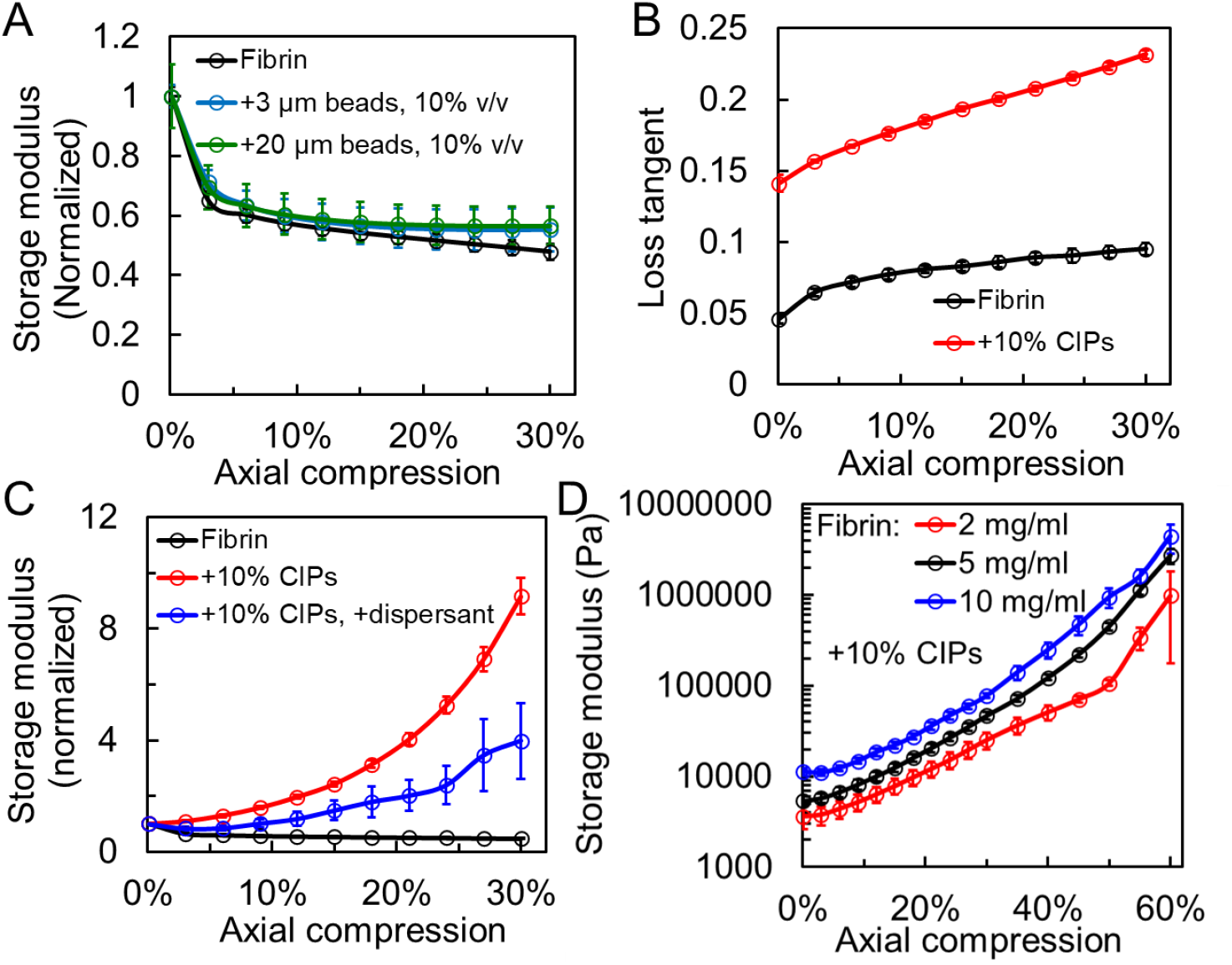
Mechanistic investigation of compression stiffening in fibrin gels: size effects, jamming, particle aggregation, and inclusion-network interaction. **A.** Normalized shear storage modulus as a function of compressive strain for fibrin network embedded with 3 μm or 20 μm monodisperse PS beads. **B**. Loss tangents as a function of axial compressive strain for fibrin hydrogels with and without 10% v/v CIPs. **C**. Normalized shear storage modulus of fibrin gels with and without the dispersant under axial compressions. **D**. Shear storage modulus as a function of compressive strain for fibrin gels containing 10% v/v CIPs with varying fibrinogen concentrations. n≥3 for all curves. Error bar denotes SEM.

We also investigate jamming as a potential explanation for the observed compression stiffening. We note that even at a compression level of 30%, the effective volume fraction of CIPs is only 10%/ (1-30%)=14.3%, well below the three-dimensional jamming thresholds (e.g., 64% for monodisperse spherical particles) (43, 44). A recent study of strain-controlled jamming in soft composites further confirms that particle contacts dominate only at considerably higher effective solid fractions (45). Moreover, the loss tangents of the fibrin exhibit a slight increase during compression (Fig. 2B). This behavior contrasts with that of a jammed system, where the loss tangent typically decreases as the particle volume fraction increases and jamming intensifies (46-48). Furthermore, since the particles are embedded within the fibrin network, the compression stiffening we observe cannot be attributed to the jamming mechanism for attractive particles, which involves the formation of fractal clusters and can reach jamming at low volume fractions (49).

We next investigate the impact of CIP aggregation. We apply a dispersant comprising 2% Tween-20 and 2% polyvinyl pyrrolidone (1300 kDa) to the fibrin pre-gel mixture. This dispersant effectively reduces CIP aggregation in the fibrin hydrogel (Fig. S9A). The average equivalent diameters of CIP aggregates decrease from 17 μm to 7.7 μm in fibrin. Correspondingly, the dispersant suppresses the compression stiffening observed in these gels (Fig. 2C and Fig. S9B). Additionally, we notice that CIP aggregation is weaker in the collagen network, which correlates with a smaller compression stiffening effect (Fig. S6). These results suggest that CIP aggregation plays an important role in the compression stiffening properties of fibrin gels. Furthermore, our investigation reveals that compression stiffening is not solely attributed to the inclusions alone, but rather involves interactions between the inclusions and the network, as alterations in fibrinogen concentration modulate the absolute magnitude of compression stiffening (Fig. 2D). This result suggests that the mechanical behavior of the hydrogel is influenced by the interplay between inclusion morphology and the fibrin network structure, a mode mimicking biological tissues where cell-matrix complex interactions impact the tissue mechanics.

The aggregation of particles in the fibrin hydrogel not only increases the effective size of the inclusions but also alters their shape. This raises the question of whether the effects of aggregation on compression stiffening arise from polydisperse size distributions or variations in morphology. To examine the impact of polydisperse inclusions, we introduce PS bead inclusions with polydisperse sizes containing 3 μm, 10 μm, and 20 μm. The quantity ratio of the beads is 1:1.6:2.2, corresponding to a ratio of volume fraction 1:4:10, which mimics the size distribution of CIP inclusions. Rheological results show that, similar to control fibrin, fibrin with polydisperse inclusions at 10% volume fraction softens during compression (Fig. 3A). Although we have only tested one quantity ratio, we do not observe any indication of compression stiffening caused by inclusion size polydispersity, suggesting that the observed compression stiffening in gels containing CIPs is not solely due to size distribution but may instead require variations in inclusion shape.

**Figure 3.**
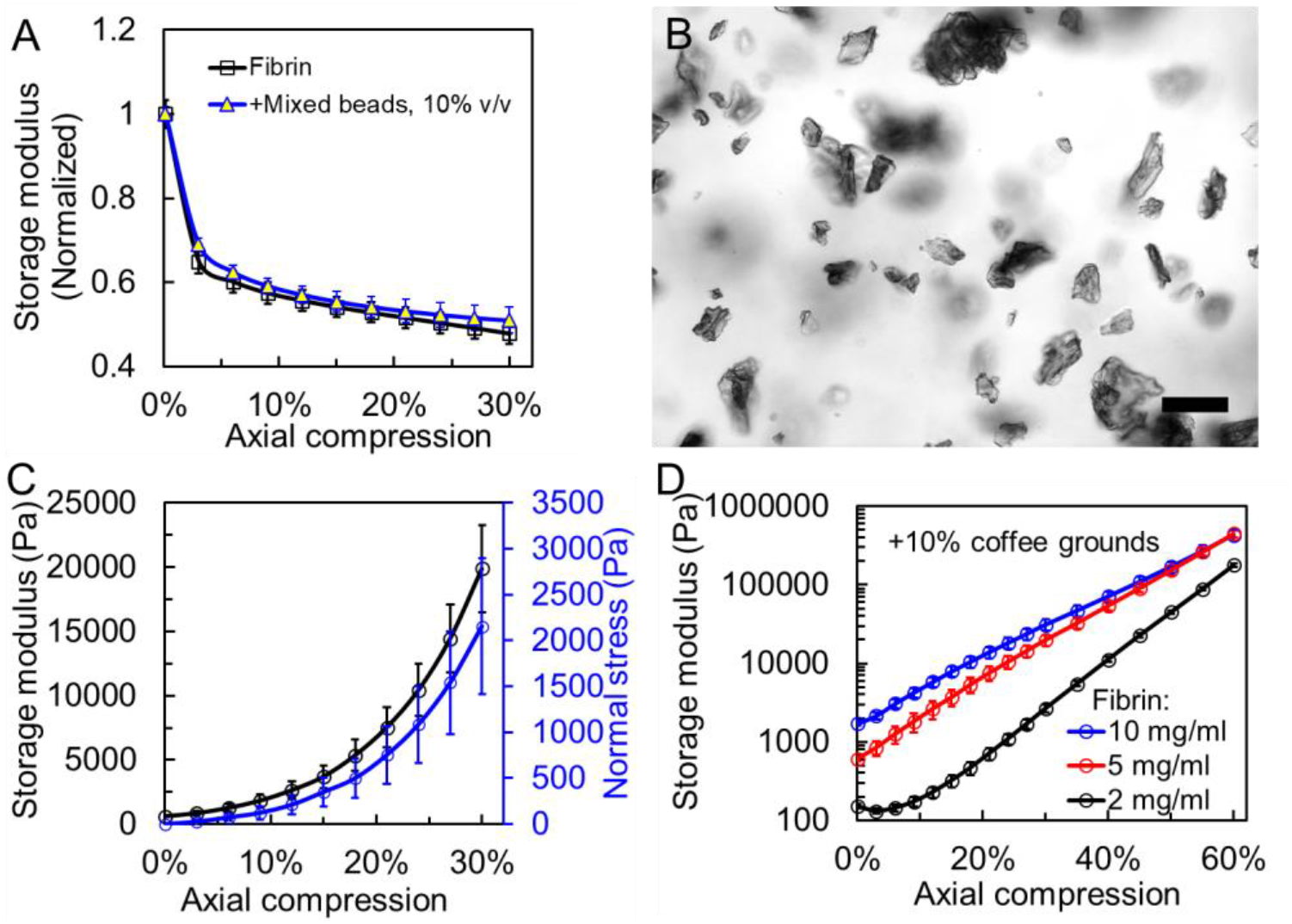
Role of inclusion size and morphology in compression stiffening. **A.** Normalized shear storage modulus as a function of compressive strain for fibrin gels embedded with polydisperse spherical PS beads. The initial modulus for fibrin containing polydisperse particles is 788 Pa. **B**. Representative bright-field image of coffee ground particles, which exhibit a wide range of sizes and diverse morphologies. The particles are retained in a three-dimensional PDMS matrix to more precisely visualize their morphologies. Scale bar = 50 μm. **C**. Shear storage modulus and axial stress as a function of axial compressive strain for fibrin gels containing 10% coffee grounds. **D**. Shear storage modulus as a function of compressive strain for fibrin gels embedded with 10% coffee ground particles at varying fibrinogen concentrations. n≥3 for all curves. Error bars denote SEM.

To test the role of inclusion morphology, we next investigate whether the compression stiffening caused by CIP aggregates can be replicated using another type of particle with diverse morphologies and similar sizes. We find that finely ground coffee particles exhibit a variety of shapes and sizes, largely ranging from 5 to 50 μm (Fig. 3B, Fig. S10). These particles are neutralized and repeatedly washed (see Methods), and the supernatant from these particles does not alter the compression softening response of the fibrin network (Fig. S11A). When embedded in fibrin gels, coffee grounds, unlike CIPs, are more uniformly distributed within the network, do not significantly form aggregates (Fig. S11B), and only moderately increase gel stiffness by 33%. Remarkably, we observe compression stiffening of approximately 33-fold in fibrin with 10% volume fraction coffee grounds upon 30% compression (Fig. 3C), replicating the effects seen with CIP inclusions. Interestingly, this fibrin gel containing coffee ground particles exhibits shear stiffening, yet axial compression transforms it into tissue-like shear strain softening (Fig. S12). Additionally, we find that the level of compression stiffening depends on fibrinogen concentration, also underlining the role of fiber-particle interactions (Fig. 3D). These findings collectively suggest that the diverse morphologies of inclusions play a significant role in driving tissue-like mechanical behavior.

We next investigate how the diverse morphologies of inclusions contribute to compression stiffening. We first refer to the fibers stretching around stiff spherical inclusions mechanism (23, 24). In this mechanism, each fibrin filament is modeled as a rope-like soft spring. Compression of the composite network containing spherical particles leads to particle contact percolation, which depends on both the initial particle volume fraction and the degree of compression. Because the particles are stiff and thus non-deformable, their displacements upon reaching or surpassing the percolation threshold become strongly non-affine. With further compression, this nonaffinity of particle motion shows a pronounced increase, reflecting strongly heterogeneous particle displacements. In particular, lateral rearrangements of contacting particles pull neighboring fibrin filaments into extension, generating localized tensile strain. Due to the rope-like elasticity of fibrin filaments, these tensile stresses can propagate through the network, forming a tension-dominated stress pathway that results in macroscopic stiffening. Notably, this stretching-dominated stress propagation occurs prior to and independently of jamming.

A key factor in this mechanism is the occurrence of contact percolation, which is determined by both the initial particle volume fraction and compression level. We therefore hypothesize that inclusions with irregular shapes (CIP aggregates and coffee grounds) have a lower percolation threshold than spheres (which we find to be ∼16% in 3D), allowing compression stiffening to occur at much lower volume fractions. Our numerical simulations support this hypothesis, showing that even prolate and oblate ellipsoid inclusions percolate at lower thresholds than spheres (Fig. 4 and Supplementary Note II). Additionally, the CIP aggregates and coffee ground particles present larger contact areas and likely exhibit greater interparticle friction compared to spheres. These factors together enhance the stability of the percolated structure, while the increased friction may also amplify the magnitude of non-affine particle motions (50, 51). However, simulating the full percolation dynamics of irregularly shaped particles remains a significant challenge.

**Figure 4.**
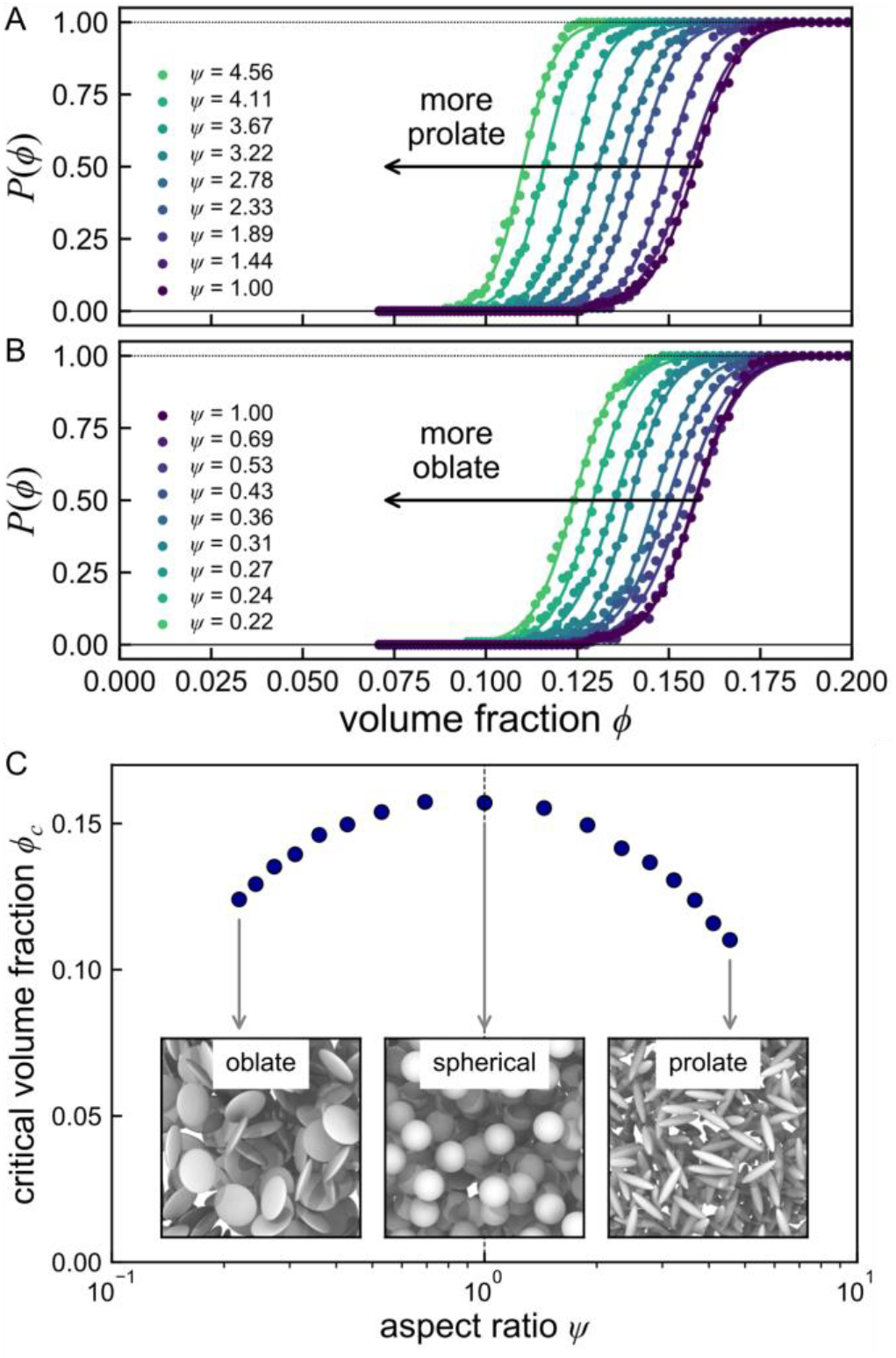
Effect of particle asphericity on the contact percolation threshold. **A** and **B.** Probability of contact percolation *P*(*ϕ*) as a function of the particle volume fraction *ϕ* with varying particle aspect ratio *ψ* for (A) prolate particles with *ψ* > 1 and (B) oblate particles with *ψ* < 1. Markers correspond to simulation results, averaged over 100 samples, and solid lines correspond to fits to a sigmoid *P*(*ϕ*) = (1 + *k* exp(*ϕ* − *ϕ*_*c*_))^−1^. **C**. Critical volume fraction *ϕ*_*c*_ for contact percolation as a function of the ellipsoid aspect ratio Ψ. The critical volume fraction Φ_*c*_ exhibits a maximum when particles are spherical (Ψ = 1), decreasing as particles become either increasingly oblate (Ψ < 1) or prolate (Ψ > 1). Inset images depict particles with aspect ratios indicated by arrows.

To study particle percolation within the fibrin network, we conducted three sets of experiments to test the hypothesis above. First, we note that the percolation of solid inclusions alters the composite’s material properties, such as its modulus (52-55). We therefore anticipate that once percolation is reached, increasing the density of percolated particles by compression will lead to stiffening of the composite material, independent of the polymer network properties. We test this in polyacrylamide (pAAm) and agarose hydrogels. We observe that CIPs form aggregates within the pAAm gel (Fig. S13A). Interestingly, the linear pAAm gel exhibits compression stiffening when embedded with 10% v/v CIPs (Fig. 5A, Fig. S13B), although the absolute stiffening upon 20% compression in pAAm (1.8 kPa) is much smaller than in fibrin (15 kPa). This stiffening effect is removed when a dispersant is used to reduce CIP aggregation (Fig. 5A, Fig. S13B and C), further confirming that particle aggregation, rather than volume fraction alone, drives compression stiffening. Similarly, coffee ground particles transform the pAAm gel from a linearly elastic material to a compression-stiffening one, whereas polydisperse spherical PS beads do not (Fig. S14A and B). Additionally, coffee ground particles convert agarose, a hydrogel composed of linear polysaccharide, from compression softening to compression stiffening (Fig. S14C).

**Figure 5.**
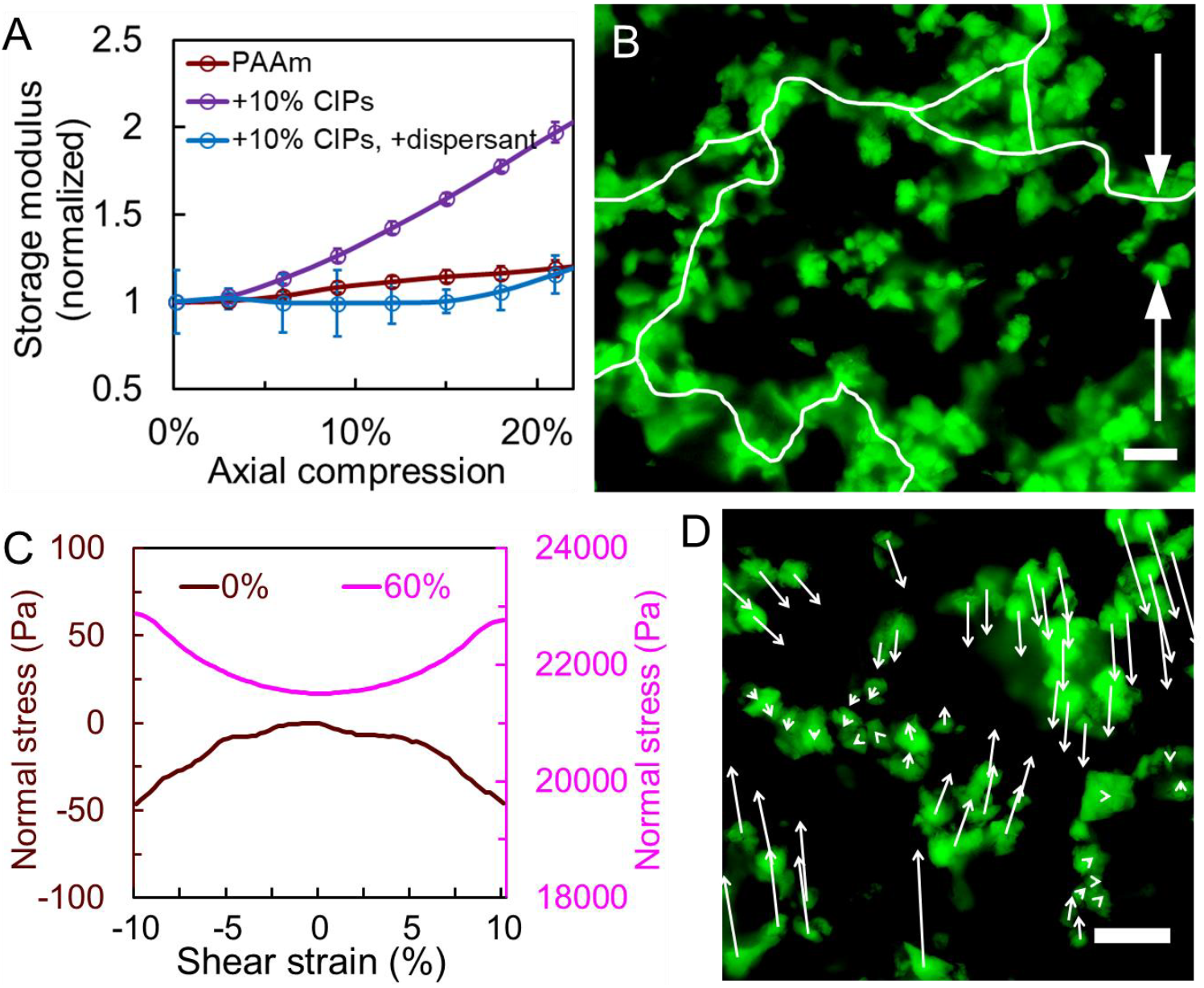
Percolation network and its role in compression stiffening. **A.** Normalized shear storage modulus of pAAm gels embedded with 10% v/v CIPs, with and without dispersant, as a function of axial compressive strain. **B**. Imaging of the percolation path formed by coffee ground particles (in green) in fibrin gels containing 10% coffee grounds upon 30% compression. The percolation path is indicated with the white curve, and the compression direction is along the y-axis, as indicated by white arrows. **C**. Normal stress as a function of shear strain for fibrin gels embedded with 10% coffee grounds, with or without 60% axial compression. **D**. Non-affine displacement of coffee ground particles in fibrin gels containing 10% coffee grounds upon 30% compression. The image shows coffee particles (green) prior to compression, with the displacement vectors of individual particles after 30% compression indicated by white arrows. The compression direction is the y-axis, as in B. n≥3 for all curves. Error bars denote SEM. Scale bar = 50 μm.

Overall, we observe that irregularly shaped inclusions induce compression stiffening in multiple hydrogel systems. Similar trends have been reported in PDMS/glass bead composites, confirming the role of rigid inclusions in altering composite properties (56). However, since particle-particle contacts alone can enhance composite stiffness, even without the formation of a fully percolated particle network, by affecting stress transfer through the composite (52, 57), these rheological results support the idea that CIP aggregates or coffee ground particles at 10% volume fraction form interparticle contacts or local networks that alter local deformation. Nevertheless, they do not provide definitive evidence for the formation of a system-spanning percolated network, which requires additional evidence.

Therefore, and secondly, we directly image the percolated network of coffee grounds in fibrin after 30% compression. At this level of compression, the effective particle volume fraction increases to 10%/ (1-30%)=14.3%. Due to the opacity of coffee particles, we only image the gel surface with an imaging depth of approximately 30 μm and capture a percolation path (Fig. 5B). Notably, because percolation is inherently three-dimensional, the probability of detecting a percolation path in a given image section is relatively low; in our experimental case, it is approximately 5%. Thirdly, since a percolated particle network can withstand compressive forces but not tensile forces (see Supplementary Note III), we expect positive normal stress under large shear strain. Indeed, we observe that fibrin mixed with coffee ground particles undergoes a transition from a negative Poynting effect (58) to a positive Poynting effect (Fig. 5C, Fig. S15). Together, these experiments provide direct structural and mechanical evidence that particles with irregular shapes can form a percolated network under compression. This network contributes to mechanical resistance in the composite and may underlie the radical compression stiffening response observed in fibrin gels.

We next show that during compression, particle contact, even prior to full contact percolation, induces non-affine displacements of the inclusions (Fig. 5D, SI video 1). We also observe rotational motion of individual particles, which is likely due to high friction between irregularly shaped contacting particles and their resistance to sliding (SI Video 1). These non-affine motions lead to localized stretching of fibrin fibers, occurring either before or concurrently with contact percolation, and can be categorized into three representative modes: (1) dislocation between particle clusters, (2) rotation of individual particles, and (3) collapse of particle clusters (Fig. S16, SI Videos 2-4). This localized fiber stretching, driven by non-affine particle dynamics, ultimately contributes to gel stiffening through the fiber-stretching-around-stiff-spherical-inclusions mechanism (23).

The percolated particle networks in fibrin gels also explain the unusually high shear modulus observed under large compressions (Fig. 2D and 3D). For example, at 60% compression, the shear modulus of 5 mg/ml fibrin containing 10% coffee particles reaches approximately 0.4 MPa (Fig. 3D). This value cannot be explained solely by the stiffness of a single fibrin fiber (59-61), which has a Young’s modulus of around 10 MPa, given its effective volume fraction (∼2%). However, since coffee particles are much stiffer (∼90 MPa) than single fibrin fibers and have a larger volume fraction, and given that the radial expansion of the gel during compression is minimal (Fig. S17) where compression-induced percolation density is enhanced compared to volume-conserving materials, the mechanical resistance provided by the percolated particles can account for the observed high shear modulus. Notably, the compression stiffening curves we obtain are relatively smooth; under increasing compression, we do not observe a sharp transition between a regime in which the overall stiffness is dominated by the stretching resistance of fibers and one in which it is clearly dominated by the particle stiffness. These observations are qualitatively consistent with prior simulation results (23), where similarly smooth compression-stiffening curves were obtained for inclusion-filled networks in which the particle stiffness greatly exceeded the fiber stiffness, even as compression drove the particles beyond the jamming threshold. Together, our results and analyses show that the compression stiffening of fibrin gels containing inclusions with diverse shapes arises from both the percolated network of inclusions and the resulting tension-dominated stress propagation in fibrin fibers.

In conclusion, we demonstrate a novel mode of compression stiffening in biopolymer networks induced by the inclusion of aggregated and irregularly shaped particles, which produce significant stiffening even at low volume fractions. Our results highlight that inclusion morphology, rather than polydispersity alone, plays a critical role in achieving tissue-like mechanical properties in hydrogel composites. We show that this behavior arises from the formation of a percolated inclusion network, which forms at relatively low volume fractions due to the morphological features of the inclusions. This percolated structure facilitates tension-dominated stress propagation in fibrin fibers, resulting in compression stiffening. Additionally, the percolated inclusion network provides mechanical reinforcement under compression, further contributing to the observed stiffening response. Together, these findings advance our understanding of how the morphological characteristics of tissue components influence tissue mechanics and offer design principles for engineering biomaterials with physiologically relevant mechanical responses, with potential applications in tissue engineering and regenerative medicine.

## Materials and Methods

### Fibrin hydrogels

Fibrinogen was isolated from bovine plasma following a previously established protocol (62). Thrombin, isolated from salmon plasma (Salmonics Inc., Brunswick ME), was diluted in PBS containing 0.1% BSA to a concentration of 1000 Units per milliliter (U/ml). Fibrin gels, with or without particle inclusions, were prepared using 5 mg/ml fibrinogen, 3.3 U/ml thrombin, and 2 mM Ca^2+^ in Tris buffer (50 mM Tris, 150 mM NaCl, pH 7.4), unless specified otherwise. All fibrin gels were formed at room temperature. Pre-gel solutions were placed between rheometer plates and allowed to polymerize for 1.5 to 2 hours before rheological measurements, unless stated otherwise. The gels were surrounded with Tris buffer to prevent drying.

### Fine coffee ground particles

Fine coffee ground particles were purified from everyday-use espresso grounds. After extracting espresso, the coffee grounds were dissolved in water and neutralized to pH 7.4. Subsequently, the grounds were fully mixed in the solution again and allowed to precipitate for about 1 minute. The supernatant was poured off for further purification, while the precipitate was discarded. Then, the top fraction underwent extensive purification via repeated centrifugation at 1200 x g RCF and dissolving in water. Finally, the fine coffee grounds were dried in an oven at 37°C. The volume fraction was estimated based on the assumption that the coffee grounds have the same density as water (1 g/cm^3^) and do not swell when dissolved. Therefore, the weight per volume (w/v) concentration was used to represent the volume fraction. The stiffness of the coffee particles was assumed to be the same as the stiffness of the coffee beans from which they were derived. The compressive modulus was determined by compressing coffee bean segments with flat ends, yielding a measured modulus of 86 MPa.

### Fibrin gels with particle inclusions

Except for CIPs, particles were mixed with the pre-gel solution to form fibrin gels embedded with particles at specified volume fractions. Polystyrene beads (3 μm, 10 μm, and 20 μm, Sigma Aldrich) originally had a volume fraction of 10% upon purchase and were concentrated via centrifugation before mixing with the fibrin pre-gel solution. All hydrogels were fabricated to a thickness of 1 mm. To prevent particle precipitation and ensure rapid gelation, fibrin gels containing coffee grounds employed 5 U/ml thrombin, resulting in gelation within 30 seconds.

Fibrin gels with CIP inclusions were synthesized vertically in a customized chamber to eliminate particle precipitation-induced inhomogeneity (Fig. S18A), and the homogeneity of the gel was verified (Fig. S18B and C). The diameter of CIPs (Sigma Aldrich, manufactured by BASF, Germany) ranged from 2-8 μm. The volume fraction of the CIP solution was determined by the difference in solution volume before and after CIP dissolution, assuming no swelling and no water absorption by the CIPs. For adequate mixing, fibrin gels containing CIPs employed 1 U/ml thrombin for 5 and 10 mg/ml fibrinogen formulations, and 2.5 U/ml thrombin for 2 mg/ml fibrinogen formulation. The pre-gel mixture was injected into the glass chamber 60-70 seconds after adding thrombin to initiate gelation. To image and illustrate CIP aggregation inside the fibrin gel, CIPs were used at 1% v/v, as they appeared completely dark under bright field imaging. For fibrin gels containing dispersant (2% Tween-20 and 2% polyvinyl pyrrolidone, 1300 kDa), the dispersant slowed fibrin polymerization and gelation, and thus thrombin was increased to 7 U/ml. After full gelation, one coverslip was removed, and the fibrin gel was cut into an 8 mm diameter cylinder for rheology measurements. To minimize slipping between the fibrin gel and the rheometer plates, the top and bottom of the gel were glued. When attaching the gel to the rheometer, the normal force sensor of the rheometer was carefully monitored to ensure that the axial stress on the top surface was zero, serving as the 0% compression level. Disturbance to the fibrin gels during the transfer process was minimal, as the gel thickness measured by the rheometer remained approximately 1 mm.

For imaging, fibrin gels containing CIPs or coffee ground particles were also synthesized in the customized chamber. Gels were imaged without removal from the chamber. To apply horizontal compression during imaging, the glass slide on one side of the gel was manually displaced toward the other side by a distance corresponding to the required compression level. In some experiments, red fluorescent nanoparticles (200 nm, Carboxylate-Modified, Thermo Fisher) were added at 0.015% v/v to visualize the fibrin network. Additionally, 8 μg/ml fluorescein was used to stain the aqueous phase, allowing coffee particles to be imaged via fluorescence microscopy since they appear dark and lack fluorescence, and their fluorescence images are presented with an inverted color.

### Polyacrylamide hydrogels

Similar to the preparation of fibrin gels embedded with CIPs, polyacrylamide (pAAm) gels were synthesized vertically using an SDS-PAGE gel caster with a 1 mm thickness, adapted from a previous approach (20). Pre-gel solutions contained PBS, 3.6% acrylamide, and 0.25% bis-acrylamide, with or without CIPs and the dispersant. Polymerization was initiated with 0.3% APS and 0.25% TEMED, after which the mixture was injected into the gel caster to allow vertical gelation. Only the center region of the hydrogel was used for further measurements. The gel was cut into an 8 mm diameter and glued between the rheometer plates for rheological measurements.

### Rheometry

The mechanical properties of the hydrogels were determined using a stress-controlled Kinexus rheometer equipped with an 8 mm diameter geometry plate. Shear modulus was measured with 1% shear strain at 1 Hz. Axial strains were applied by adjusting the gap size between the geometry plate and the bottom plate during shear oscillation (Fig. S5). Compression was applied stepwise in increments of 3% axial strain up to a total of 30%, and in some experiments, further in 5% increments up to 60% total compression, unless specified otherwise. Following each compression step, the shear modulus typically exhibited an instantaneous increase followed by relaxation, and the next compression was applied only after complete relaxation. All measurements were conducted at room temperature.

## Acknowledgement

The work of XS and PAJ was supported by grants DMR-2309043 and CMMI-1548571 from the US National Science Foundation, R35-GM-136259 from the National Institute of General Medicine Sciences, and R01-EB-017753 from the National Institute of Biomedical Imaging and Bioengineering. PAJ thanks Corey O’Hern for helpful discussions. JLS acknowledges support from the Eric and Wendy Schmidt AI in Science Postdoctoral Fellowship, a Schmidt Sciences program.

The authors declare no conflict of interest.

