## Supplementary Information for "Tissue-like compression stiffening in biopolymer networks induced by aggregated and irregularly shaped inclusions"

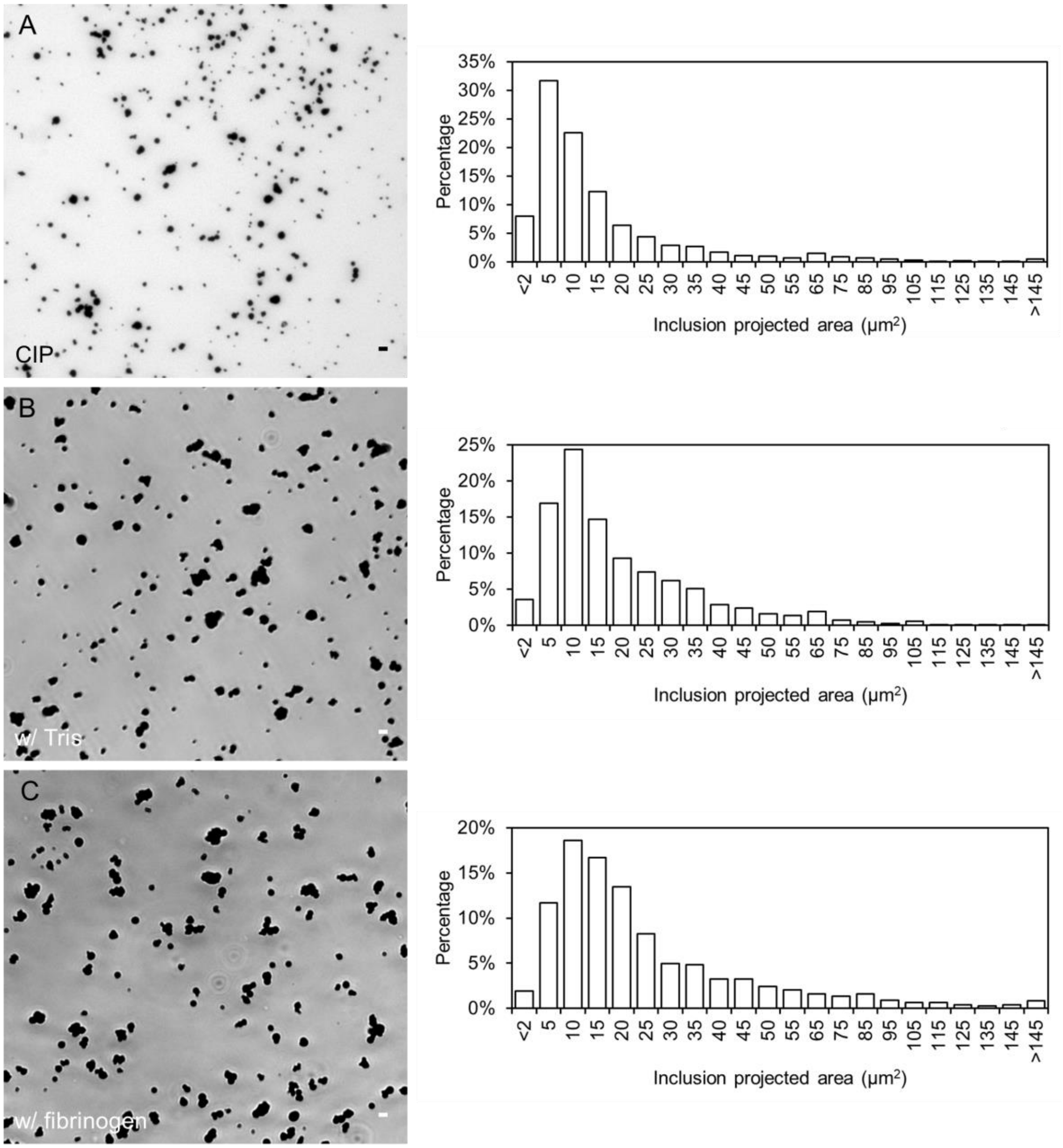


**Figure S1**. CIP morphologies and aggregation. **A**. Native CIPs before dissolution. **B**. CIPs dissolved and precipitated in Tris buffer. **C**. CIPs dissolved and precipitated in fibrinogen solution. Scale bar=5 μm. Right of each image: Size distribution of CIPs under each condition. Number of inclusions analyzed n>800.

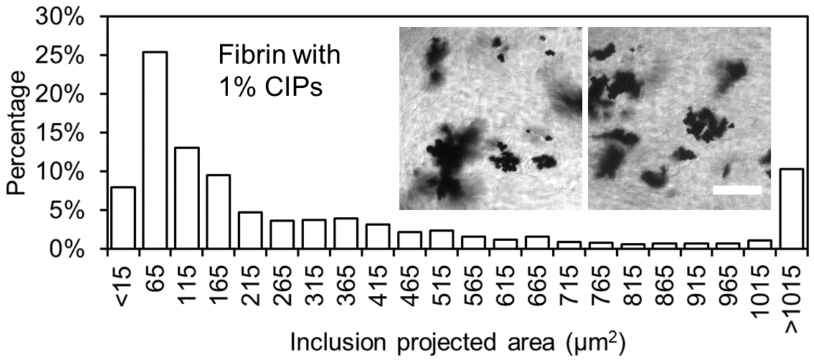


**Figure S2**. Size distribution of CIP inclusions in fibrin hydrogel. Two-dimensional projected images were acquired, and the projected area of each aggregate was analyzed. Insert: Bright field images showing representative CIP aggregates in fibrin. The volume fraction of CIPs is 1% instead of 10% to conduct bright field imaging. CIPs spontaneously form aggregates during fibrin polymerization. Scale bar=50 μm. Number of inclusions analyzed n>800.

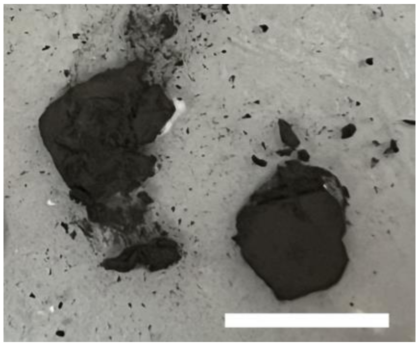

**Figure S3**. Rusting examination of CIPs inside fibrin gels. Fibrin gels were embedded with 10% v/v of CIPs. The CIPs remain stable and rust-free for at least two months, even after the fibrin gels are partially hydrolyzed. Scale bar=10 mm.

**Supplementary Note I**. Enhanced stiffness of fibrin by CIP inclusions.

We observe that the inclusion of CIPs in fibrin gels at only 10% volume fraction leads to a 10-fold increase in stiffness, even without applied compression. While the primary focus of this study is to systematically investigate compression stiffening induced by particle inclusions, this reinforcement effect suggests a potential mechanism contributing to compression stiffening. Therefore, we aim to identify the origin of the enhanced stiffness in the absence of compression.

A previously proposed mechanism (1) demonstrates that a 10-fold increase in stiffness at a 10% volume fraction can arise from attractive interactions between inclusions and the hydrogel matrix. These interactions induce localized polymer densification, effectively increasing the filler's effective size and facilitating the formation of a percolated network, which enhances the overall stiffness of the hydrogel. The key element of this mechanism is the attractive filler-matrix interactions, which promote local matrix densification around inclusions without requiring particle aggregation.

We show that CIPs exhibit high affinities for both unpolymerized fibrinogen and polymerized fibrin fibers (Fig. S4), supporting the existence of a matrix-densification-around-inclusions mechanism in our system. To further assess the role of CIP aggregation, we introduce a dispersant (2% Tween-20 and 2% polyvinyl pyrrolidone) to reduce CIP aggregation (Fig. S9A). Although Tween-20 is a nonionic detergent, we find, unexpectedly, that this dispersant does not weaken the CIP-fibrin attraction. Instead, reduced aggregation slightly enhances the attraction, likely due to better CIP dispersion and increased CIP-polymer contact area. Notably, fibrin gels containing CIPs with or without dispersant exhibit a similar level of stiffness enhancement (Fig. S9B), indicating that CIP aggregation does not contribute to gel reinforcement and further supporting the matrix-densification-around-inclusions mechanism. Furthermore, since fibrin gels containing CIPs with or without dispersant exhibit similar stiffness reinforcement but different compression stiffening behavior, this suggests that the matrix-densification-around-inclusions mechanism does not contribute to compression stiffening. As a negative control, we show that collagen networks exhibit weaker attraction to CIPs (Fig. S4), and correspondingly, the reinforcement effect from CIP inclusion without compression is minimal (Fig. S6). This finding further supports the conclusion that CIP-biopolymer attraction governs the stiffness enhancement of the biopolymer network, whereas compression stiffening arises from a distinct mechanism.

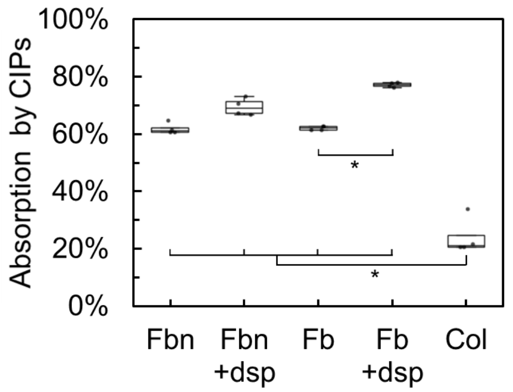


**Figure S4**. Attraction between CIPs and biopolymers, measured by quantification of protein absorption by CIPs. “Fbn”: fibrinogen at 0.1 mg/ml. “dsp”: dispersant, which contains 2% Tween-20 and 2% polyvinyl pyrrolidone. “Fb”: thrombin-polymerized fibrin at 0.1 mg/ml. “Col”: polymerized collagen formed by neutralization at 0.1 mg/ml. All proteins were used at 0.1 mg/ml to minimize network formation while maintaining the polymerization process.CIPs at 0.3% volume fraction were added to each protein solution. For polymerized fibrin and collagen, CIPs were introduced at the start of polymerization, and the mixtures were subjected to short agitation. The protein absorption level by CIPs was determined by measuring the remaining protein concentration in the supernatant via UV absorption after centrifugation and CIP removal. *$p<0.0001$.

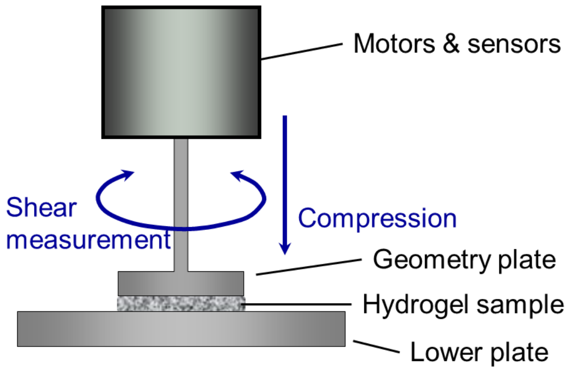


**Figure S5**. Schematic of the rheometry setup illustrating the application of axial strains to the hydrogel during shear oscillations.

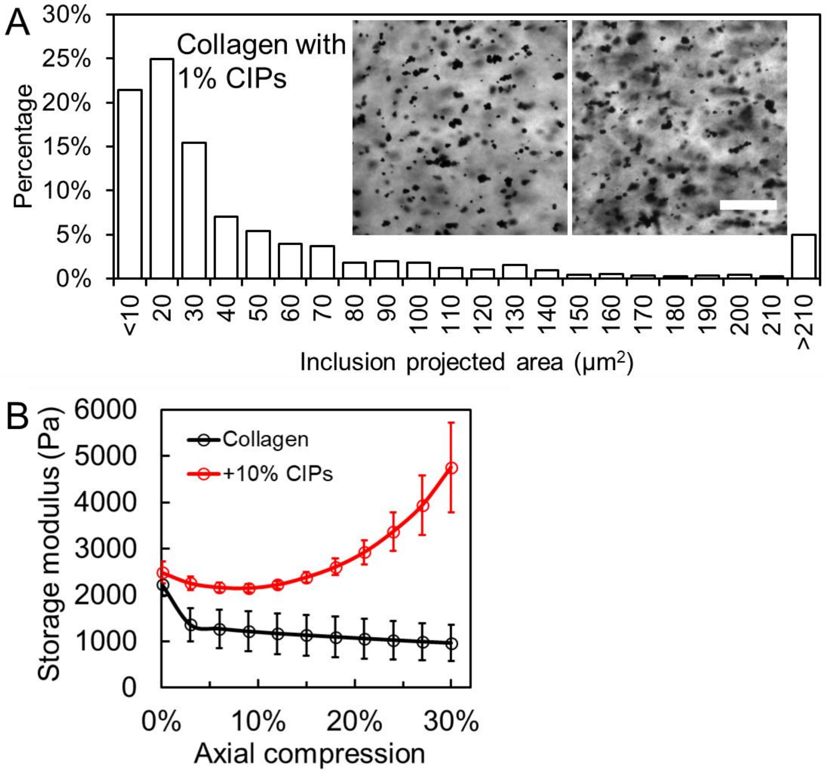


**Figure S6**. Compression stiffening in collagen networks induced by CIPs. **A**. Size distribution of CIP inclusions in collagen hydrogels. Inset: Bright-field images showing CIP aggregates in collagen. The volume fraction of CIPs is 1% instead of 10% to conduct bright field imaging. While CIPs spontaneously form aggregates, the aggregation level is lower than in fibrin networks. Scale bar=50 μm. Number of inclusions analyzed n>800. **B**. Shear storage modulus as a function of compressive strain in collagen gels with 0% and 10% v/v CIPs. n≥3 for all curves. Error bar denotes SEM.

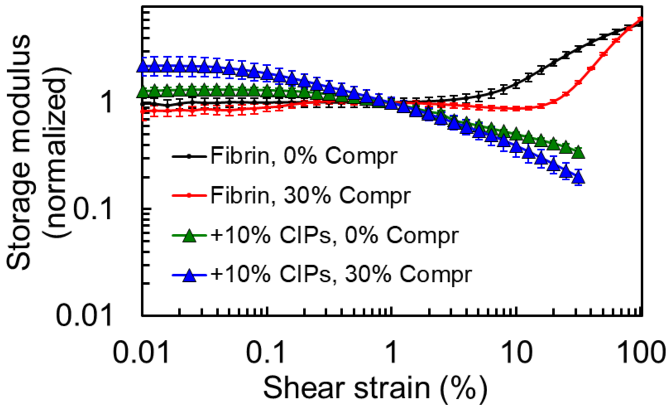


**Figure S7**. Shear storage modulus as a function of shear strain in fibrin gels containing 0% or 10% CIPs, with or without 30% axial compression. n≥3 for all curves. Error bar denotes SEM. Fibrin hydrogels are shear strain stiffening, whereas fibrin gels embedded with CIPs exhibit tissue-like shear strain softening.

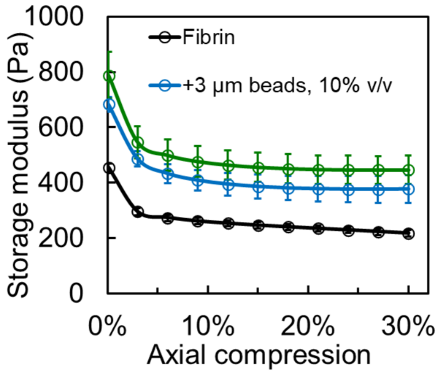


**Figure S8**. Shear storage modulus as a function of compressive strain for fibrin gels embedded with 3 μm or 20 μm monodisperse polystyrene beads. n≥3 for all curves. Error bar denotes SEM.

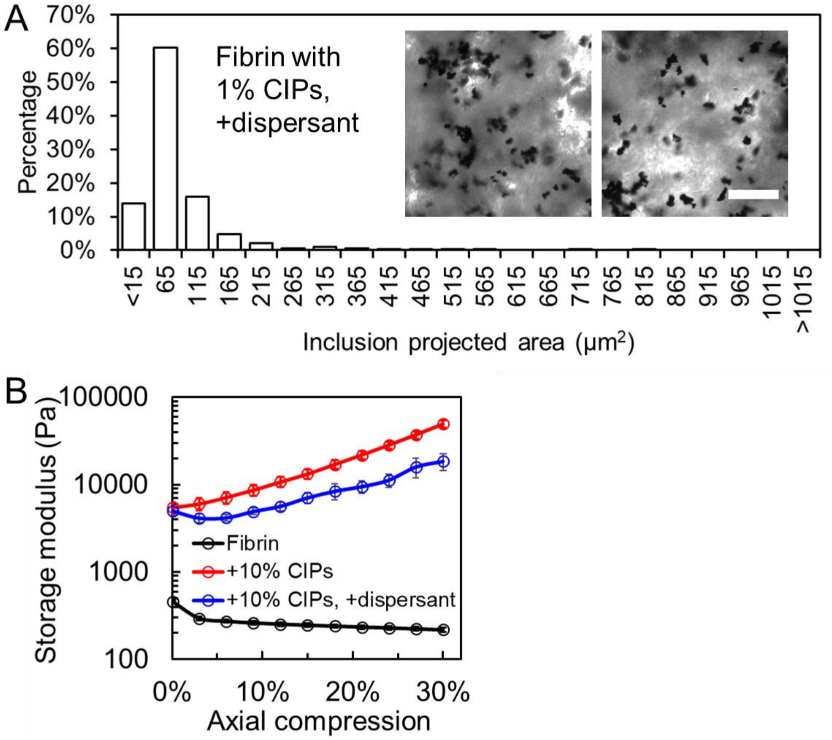


**Figure S9**. Effect of dispersant on CIP aggregation and CIP-induced compression stiffening. **A**. Size distribution of CIP inclusions in fibrin hydrogels with the dispersant comprising 2% Tween-20 and 2% 1300 kDa polyvinyl pyrrolidone. Insets: Bright field images showing CIPs in fibrin containing dispersant. The volume fraction of CIPs is 1% instead of 10% to conduct bright field imaging. Scale bar = 50 μm. Number of inclusions analyzed n>800. **B**. Shear storage modulus of fibrin gels with and without dispersant under different axial compressions. n≥3 for all curves. Error bar denotes SEM.

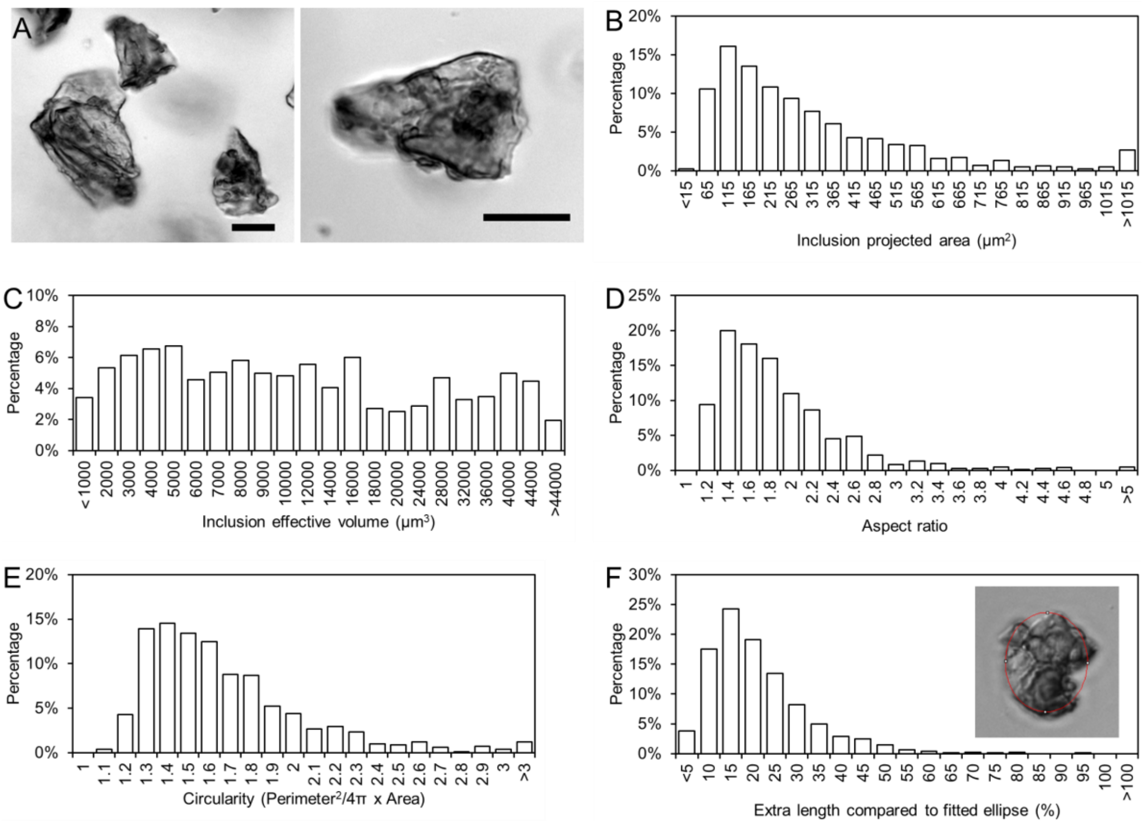


**Figure S10**. Morphological characterization of coffee ground particles. **A**. Representative images of coffee ground particles, showing concave and convex regions with multiple facets. Scale bar = 20 μm. **B-F**. Distributions of key morphological parameters: 2D projected area (B), estimated volume calculated from the projected area (C), aspect ratio (D), circularity (E), and relative extra length (F). The "extra length" in F represents the relative difference between the 2D projected perimeter of the particle and its fitted ellipse, indicating that the particle surfaces are rough with broken lines rather than smooth. Number of inclusions analyzed n>800. To ensure accurate morphological analysis, 1% coffee ground particles are embedded in a three-dimensional PDMS matrix prior to imaging, so that each particle remains fully suspended in 3D space, which prevents particles from resting flat against a surface and allows their full geometry to be captured in 2D projections.

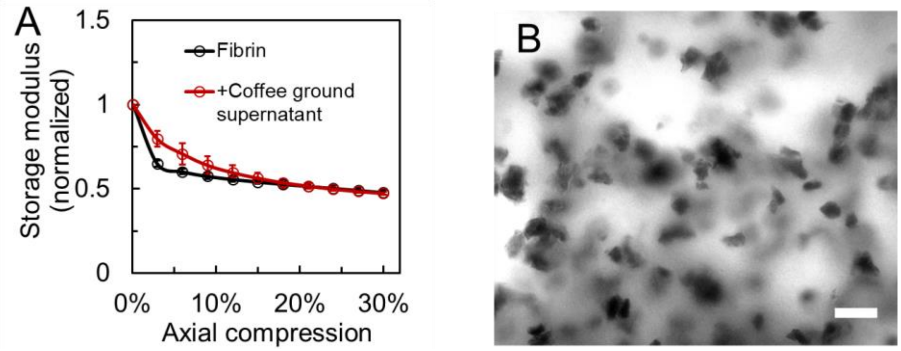


**Figure S11**. Fibrin network embedded with coffee ground particles. **A**. Normalized shear storage modulus of fibrin gels containing only the supernatant portion of coffee ground particles under compression, compared with pure fibrin. The results indicate that the supernatant does not alter the compressive response of the fibrin network. n≥3 for all curves. Error bar denotes SEM. **B**. Representative image of fibrin gel embedded with 10% coffee ground particles. Scale bar = 50 µm.

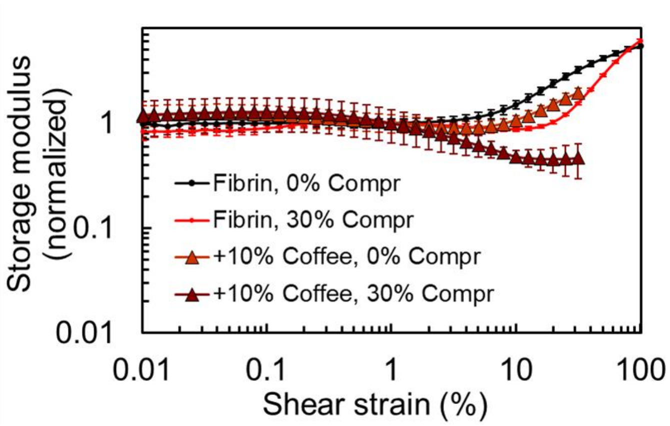


**Figure S12**. Shear storage modulus as a function of shear strain in fibrin gels containing 0% or 10% coffee ground particles, with or without 30% axial compression. Fibrin hydrogels exhibit shear strain stiffening, whereas axial compression transitions fibrin gels embedded with coffee particles from shear strain stiffening to tissue-like shear strain softening. n≥3 for all curves. Error bar denotes SEM.

**Supplementary Note II**. Mechanisms for the reduction of the contact percolation threshold

#### 1. Reduction in the contact percolation threshold due to particle asphericity

To evaluate the impact of particle asphericity on the volume fraction at which contact percolation occurs, we characterize the contact percolation transition in systems consisting of axially symmetric ellipsoidal particles as a function of the particle aspect ratio $\alpha$, defined as the ratio of the particle length along the axis of symmetry, $\psi$, to the particle length along the perpendicular axes, $b = c$. For spherical particles, $\alpha = 1$. We consider both prolate (elongated) particles, with $\psi > 1$, and oblate (disk-shaped) particles, with $\psi < 1$.

#### 1.1 Ellipsoidal particle model

A convenient model for simulating interacting ellipsoids is the Gay-Berne potential, essentially a generalization of the 12-6 Lennard-Jones potential to ellipsoidal particles. The general form of the Gay-Berne potential is given by (2-4)

$$u_{\text{GB}}(\boldsymbol{r}, \boldsymbol{A}_i, \boldsymbol{A}_j) = \eta(\boldsymbol{A}_i, \boldsymbol{A}_j) \cdot \chi(\hat{\boldsymbol{r}}, \boldsymbol{A}_i, \boldsymbol{A}_j) \cdot u_r(\boldsymbol{r}, \boldsymbol{A}_i, \boldsymbol{A}_j)$$

in which the ellipsoid shape, specified by semiaxes $a_i$, $b_i$, and $c_i$, is characterized by a shape matrix $\boldsymbol{S}_i = \text{diag}(a_i, b_i, c_i)$, the relative interaction energies (well depths) are contained in a matrix $\boldsymbol{E_i} = \text{diag}(\varepsilon_{a,i}, \varepsilon_{b,i}, \varepsilon_{c,i})$, and each particle's orientation is captured by a rotation matrix $\boldsymbol{A}_i$ (3). Above, $\boldsymbol{r} = \boldsymbol{r}_j - \boldsymbol{r}_i = r\hat{\boldsymbol{r}}$ is the vector between particle centers, and the rows of the orthonormal rotation matrix $\boldsymbol{A}_i$ are the principal axis unit vectors. The energy term $u_r$ corresponds to a shifted Lennard-Jones interaction specified by the distance of closest approach $h$, the interaction radius $\sigma$, and a shift factor $\gamma$,

$$u_r(\boldsymbol{r}, \boldsymbol{A}_i, \boldsymbol{A}_j) = 4\varepsilon(\varrho^{12} - \varrho^6)$$

with

$$\varrho = \frac{\sigma}{h + \gamma\sigma}$$

The distance of closest approach $h$ is approximately given by (2, 5)

$$h = r - \left[\frac{1}{2}\hat{\boldsymbol{r}}^T \boldsymbol{G}^{-1} \hat{\boldsymbol{r}}\right]^{-1/2}$$

in which

$$\boldsymbol{G} = \boldsymbol{A}_i^T \boldsymbol{S}_i^2 \boldsymbol{A}_i + \boldsymbol{A}_j^T \boldsymbol{S}_j^2 \boldsymbol{A}_j.$$

The distance-independent terms $\eta(\boldsymbol{A}_i, \boldsymbol{A}_j)$ and $\chi(\hat{\boldsymbol{r}}, \boldsymbol{A}_i, \boldsymbol{A}_j)$ are defined as

$$\eta(\boldsymbol{A}_i, \boldsymbol{A}_j) = \left[\frac{2s_i s_j}{\text{det}(\boldsymbol{G})}\right]^{\nu/2}$$

with $s_k = [a_k b_k + c_k c_k][a_k b_k]^{1/2}$ and

$$\chi(\hat{\boldsymbol{r}}, \boldsymbol{A}_i, \boldsymbol{A}_j) = [2\hat{\boldsymbol{r}}^T \boldsymbol{B}^{-1} \hat{\boldsymbol{r}}]^{\mu}$$

with

$$\boldsymbol{B} = \boldsymbol{A}_i^T \boldsymbol{E}_i^2 \boldsymbol{A}_i + \boldsymbol{A}_j^T \boldsymbol{E}_j^2 \boldsymbol{A}_j,$$

in which $\mu$ and $\nu$ are adjustable parameters.

Here, we use the purely-repulsive WCA-type Gay-Berne potential (2, 4, 6-10), a shifted and truncated version that retains only the repulsive contribution (4, 9, 10) and is defined as

$$u_{\text{GB}}^{\text{WCA}} = \begin{cases} u_{\text{GB}} - u_{GB}^{min}, & r \le r_c \\ 0, & r > r_c \end{cases}$$

in which the cutoff distance $r_c$ is defined as

$$r_c = 2^{1/6}\sigma - \gamma\sigma + \left[\frac{1}{2}\hat{\boldsymbol{r}}^T \boldsymbol{G}^{-1}\hat{\boldsymbol{r}}\right]^{-1/2},$$

and the energy shift term is given by

$$u_{GB}^{\min} \equiv u_{\mathrm{GB}}|_{r=r_c} = -\eta \cdot \chi \cdot \varepsilon.$$

Simulations are performed using the open-source molecular dynamics simulator LAMMPS (11), using a custom implementation of the WCA-type Gay-Berne potential. The parameters used are as follows: $\gamma = \nu = \mu = 1.0$ and $\varepsilon = \sigma = \varepsilon_{i,a} = \varepsilon_{i,b} = \varepsilon_{i,c} = \varepsilon_{j,a} = \varepsilon_{j,b} = \varepsilon_{j,c} = 1.0$.

#### 1.2 Simulation procedure

We characterize the contact percolation transition using a similar approach to that used in Refs. (12-14) for spherical particles, with the important distinction that we focus on the onset of contact percolation under applied uniaxial, rather than bulk, compression. We prepare systems by placing $N$ particles with random positions and orientations in a periodic cubic box of volume $V_{\mathrm{box}} = L_x L_y L_z$, with initial box dimensions $L_x = L_y = L_z = L_0$ chosen such that the initial volume fraction is $\phi_0 = 0.001$. The particle volume fraction is defined as $\phi = NV_{\mathrm{particle}}/V_{\mathrm{box}}$, in which the individual particle volume is given by $V_{\mathrm{particle}} = \frac{1}{6}\pi abc$. First, quasistatic bulk compression is applied in small steps to yield a particle volume fraction of $\phi_i = 0.07$. Next, quasistatic *uniaxial* compression is applied along the $z$-axis, by reducing $L_z$ in small steps to yield a final particle volume fraction of $\phi_f = 0.25$.

The uniaxial compression trajectories are then analyzed to determine the volume fractions at which contact percolation occurs. Particles are defined as *in contact* if their center-to-center distance satisfies $r \leq r_c$, in which the cutoff distance $r_c$ depends on the orientations of the two particles as described above. Equivalently, we define two particles as *in contact* if their mutual repulsion energy, as defined by the WCA-like Gay-Berne potential, is nonzero. Then, for a given configuration, we cluster the particles according to their contacts. Any two particles in contact are grouped into the same cluster. Percolation is defined as occurring when a single cluster of contacting particles spans the length of the simulation box along the $z$ axis, reconnecting to itself to form an infinite cluster. In practice, we perform an ensemble of $N_{\mathrm{samples}} = 50$ simulations and, for each volume fraction $\phi$, compute the probability $P(\phi)$ of contact percolation, defined as the ratio of the number of samples exhibiting contact percolation at a volume fraction $\phi$ to the total number of samples. We then define the critical volume fraction $\phi_c$ as the volume fraction at which $P(\phi) = 0.5$.

#### 1.3 Simulation results

We plot the probability of percolation $P(\phi)$ as a function of the volume fraction $\phi$ for prolate ellipsoids (Fig. 4A) and oblate ellipsoids (Fig. 4B) with varying aspect ratio $\psi$. Circles represent simulation values, and solid lines correspond to fits to a sigmoid function $P(\phi) = (1 + k\exp(\phi - \phi_c))^{-1}$, from which we determine the critical volume fraction $\phi_c$ satisfying $P(\phi = \phi_c) = 0.5$. In both cases, the $P(\phi)$ curves shift to lower $\phi$ as the particle aspect ratio $\psi$ deviates from $1$ (spherical particles).

In Fig. 4C, we plot the critical volume fraction $\phi_c$ as a function of the particle aspect ratio $\psi$. Arrows are added to the plot to indicate spherical, oblate, and prolate. As the aspect ratio $\psi$ deviates from $1$ (spherical particles), the critical volume fraction $\phi_c$ decreases. This is true for both.

### 2. Reduction in the contact percolation threshold due to correlations in particle

**positions**

The previous simulations have focused on the impact of particle *shape* on the contact percolation threshold. We showed that for ellipsoidal particles with initially random positions and orientations, increased asphericity (quantified by the aspect ratio $\psi$) led to a reduction in the critical volume fraction $\phi_c$ for contact percolation. However, one can also reduce the contact percolation threshold by introducing correlations in the particle positions, irrespective of the particle shape.

For example, introducing attractive interactions between particles, such as depletion interactions mediated by the surrounding medium or filament-bridging effects, can lead to the formation of extended particle clusters or even a system-spanning particulate gel (15, 16). This can, in principle, occur at almost *arbitrarily* low particle volume fractions. Notably, a particulate gel structure of this sort satisfies the contact percolation criterion by definition. Hence, for a biopolymer matrix containing a spanning particulate gel structure, we would expect to observe compression stiffening caused by the same mechanism discussed in the main text: shear and extensile strain applied to the interstitial network due to the compression-induced non-affine rearrangement of particles in contact.

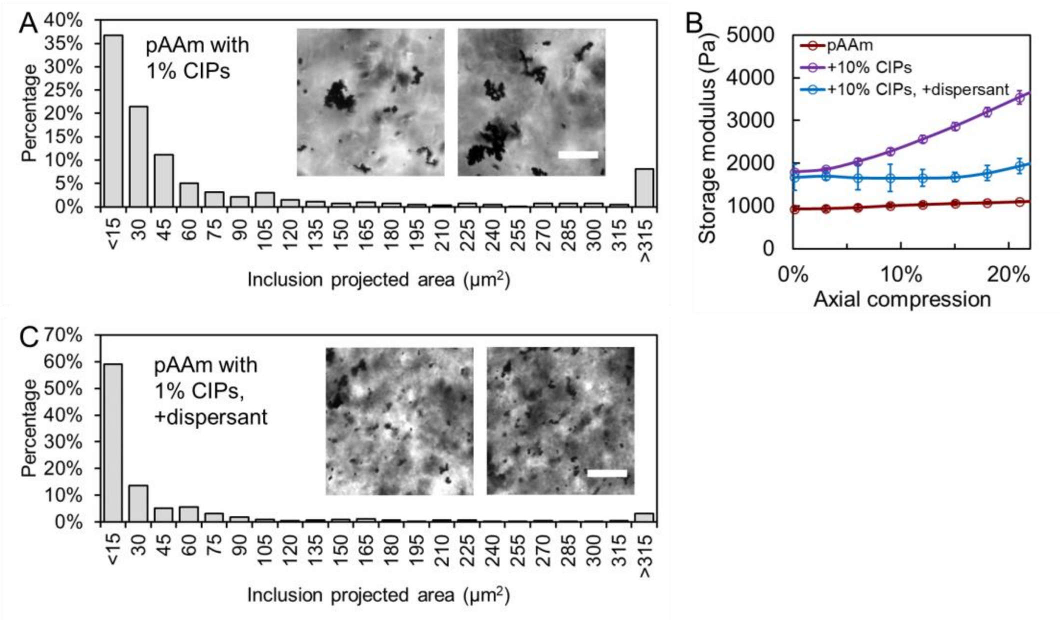


**Figure S13**. Compression stiffening in pAAm gels induced by CIPs. **A** and **C**. Size distribution of CIP inclusions in pAAm hydrogels, with (C) or without (A) the dispersant comprising 2% Tween-20 and 2% 1300 kDa polyvinyl pyrrolidone. Insets: Bright-field images showing CIP aggregation levels in hydrogels. The volume fraction of CIPs is 1% instead of 10% to facilitate bright field imaging. Scale bar = 50 μm. Number of inclusions analyzed n>800. **B**. Shear storage modulus of pAAm gels embedded with 10% v/v CIPs, with and without dispersant, as a function of axial compression. Due to the larger Poisson's ratio of pAAm gels, axial compression is accompanied by significant radial expansion, which may cause detachment from the parallel plates at large compression levels. To prevent this issue, pAAm gels were only subjected to small axial compression (~20%). n≥3 for all curves. Error bar denotes SEM.

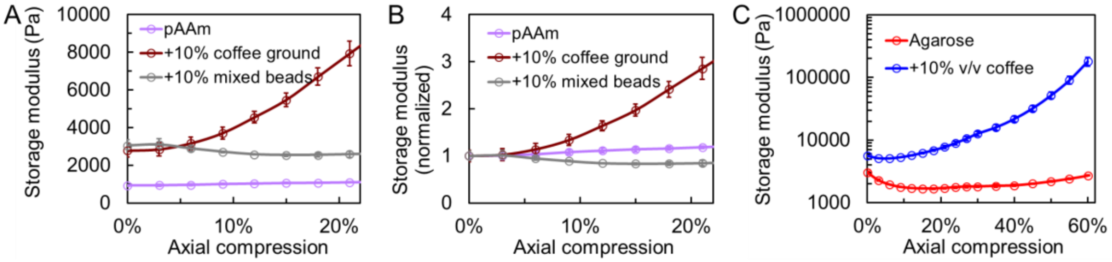


**Figure S14**. Compression stiffening in pAAm and agarose hydrogels induced by coffee ground particles. **A** and **B**. Shear storage modulus (A) and normalized shear modulus (B) of pAAm gels embedded with 10% coffee ground particles or polydisperse spherical polystyrene beads as a function of axial compression. Coffee ground particles induce compression stiffening, whereas polystyrene beads do not. **C**. Shear storage modulus of agarose gels, with or without 10% coffee ground particles embedded. n≥3 for all curves. Error bar denotes SEM.

**Supplementary Note III**. Positive and negative Poynting effects.

Most elastic materials exhibit the Poynting effect, where shearing induces a tensile stress in the normal direction to the sheared faces, causing the material to expand perpendicularly if the surfaces are not constrained. However, biopolymer networks, including fibrin hydrogels, generally exhibit a negative Poynting effect, where shear induces a compressive normal stress, pulling the plates together rather than pushing them apart. This negative normal stress arises from the intrinsic mechanical properties of semiflexible filaments that constitute the biopolymer network (17). These filaments exhibit highly nonlinear rope-like elasticity: when compressed, they buckle and become softer, whereas when stretched, they stiffen significantly. In a randomly oriented biopolymer network, an equal proportion of fibers is stretched and compressed under shear. Due to the nonlinear force-extension behavior of semiflexible filaments, stretched fibers generate larger tensile forces than compressed fibers generate compressive forces, leading to a net contractile force in the normal direction and resulting in a negative Poynting effect.

When the fibrin network is uniaxially compressed, most fibers buckle, leading to an overall softening of the gel. Under shear, even though some filaments are stretched, the majority remain buckled, which diminishes or cancels the negative Poynting effect, as observed in Fig. S15A.

In fibrin gels embedded with coffee ground particles, the majority of fibrin fibers are in a relaxed state without axial compression, allowing the negative Poynting effect to persist. However, we reckon that, when axial compression is applied, the fibrin fibers, whether stretched or compressed, cannot generate a positive Poynting effect due to their intrinsic nonlinear elasticity. On the other hand, when a percolated network of coffee ground particles forms during compression, we anticipate that a positive Poynting effect can emerge (similar to the Payne effect in rubber). Under a large shear, a portion of the percolated particle network is stretched, while another portion is compressed. These compressed structures can sustain compressive forces and exert a net pushing force in the normal direction, leading to a positive normal stress. Additionally, shear-induced compression can further enhance percolation, increasing the number of compressed particle contacts, which also contributes to positive normal stress generation. However, while some stretched structures can still support tensile forces, others may undergo partial percolation breakdown. Therefore, at large shear, percolation is expected to result in a net positive normal force.

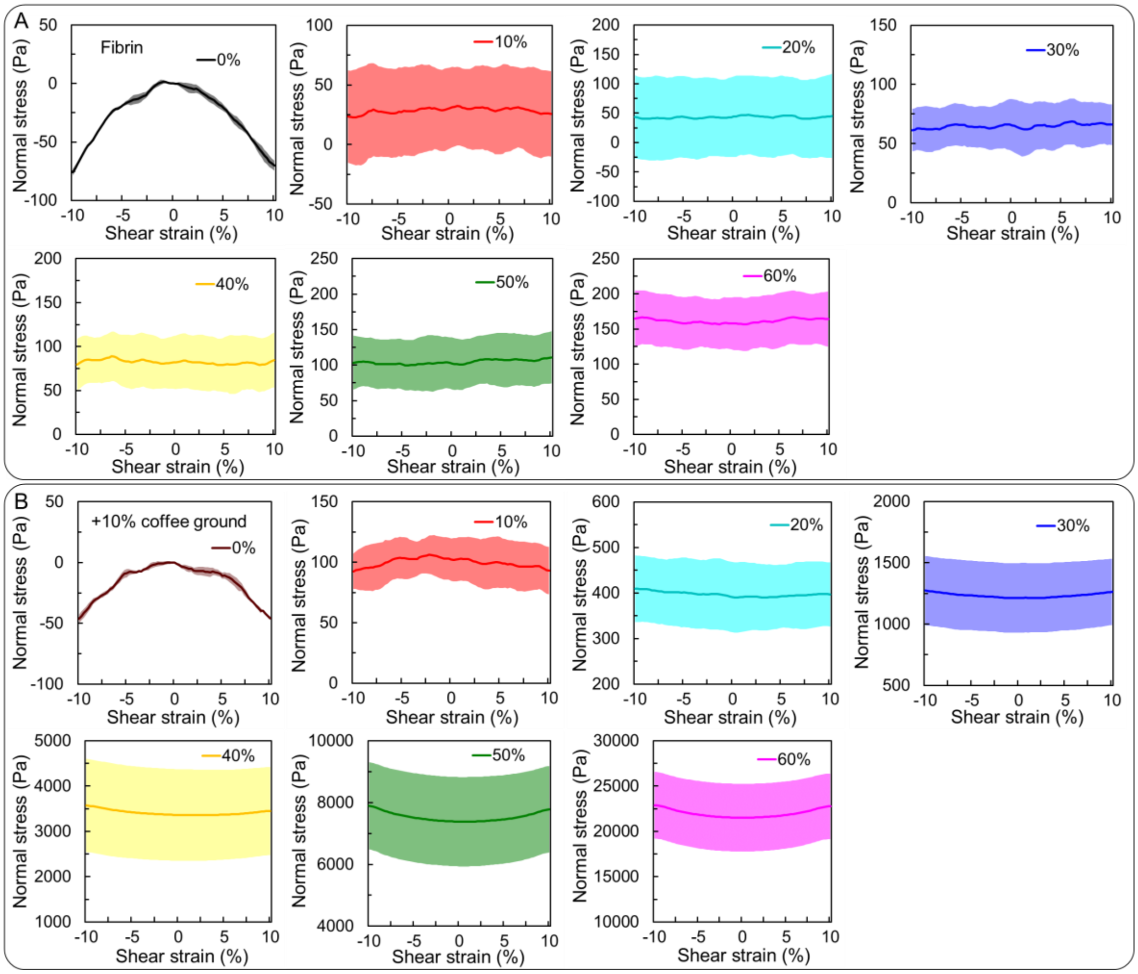


**Figure S15**. Negative and positive Poynting effect of fibrin gels. **A**. Normal stress as a function of shear strain for pure fibrin at different compression levels. Axial compression diminishes the negative Poynting effect but does not induce a positive Poynting effect. **B**. Normal stress as a function of shear strain for fibrin gels embedded with 10% coffee ground particles at different compression levels. Axial compression transitions the gel from a negative Poynting effect to a positive Poynting effect. For each condition, normal stress was allowed to equilibrate after each compression, and instantaneous normal stress was acquired during oscillations at 10% shear strain and 0.016 Hz. n≥3 for all curves. Error band denotes SEM.

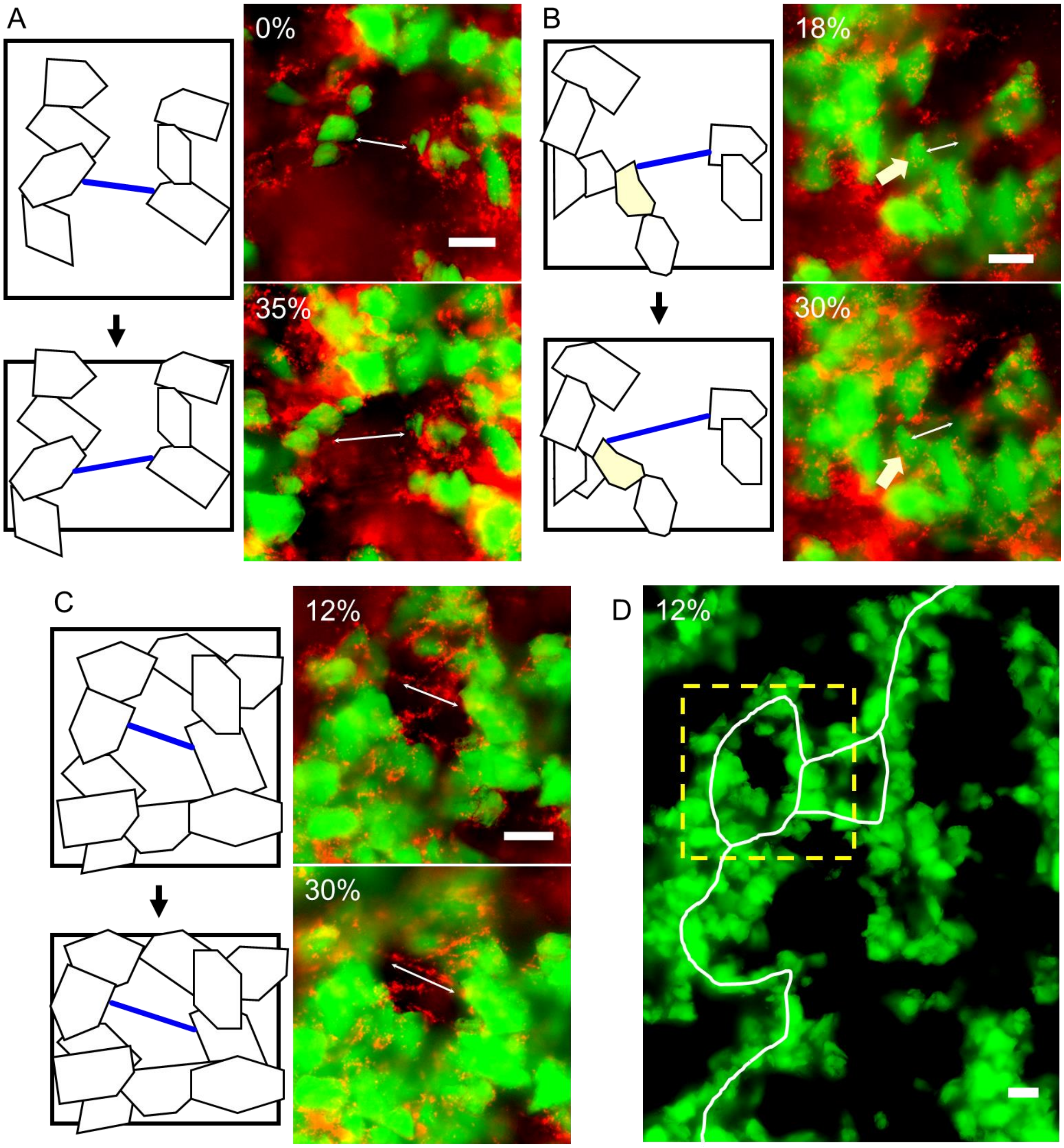


**Figure S16**. Illustrations of fibrin fiber stretching induced by non-affine displacements of coffee ground particles under compression. Three representative conditions of fibrin fiber stretching are shown in fibrin gels containing 10% coffee particles (in green) and 0.015% red fluorescent nanoparticles. While most nanoparticles are aggregated within the network, in some cases they are aligned along fibrin filaments, enabling fiber visualization. **A**. Fiber stretching due to dislocation between two particles, occurring either before or after contact percolation. **B**. Fiber stretching due to rotational motion of an individual particle (yellow arrow), occurring either before or after percolation. **C**. Fiber stretching due to the collapse of a particle cluster, occurring concurrently with contact percolation (shown in D). In A-C, the left panels show schematic diagrams before (top) and after (bottom) compression, with stretched fibers highlighted in blue. The compression direction is along the y-axis. Right panels, corresponding to SI videos 2-4, show the fluorescence images, with compression levels labeled and white arrowed lines marking the stretched fibrin filaments. **D**. Percolation map of the sample in C (in yellow box) at 12% compression. The white curve marks the percolation path. Scale bar = 25 μm.

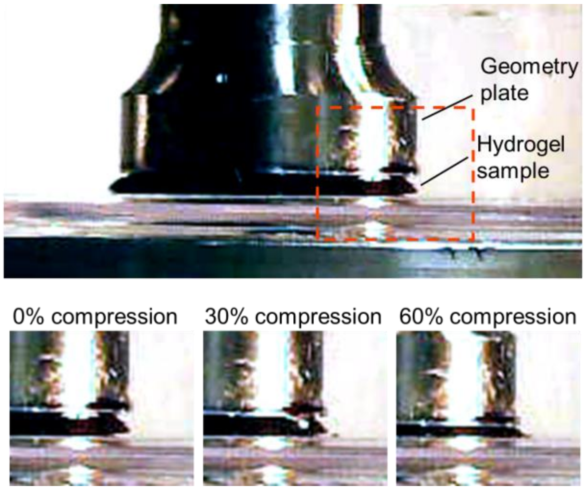


**Figure S17**. Photographs illustrating the macroscopic deformation of fibrin gels embedded with 10% coffee ground particles under axial compression. Minimal radial expansion is observed even after 60% axial compression, indicating that the gel can be approximately regarded as a fully compressible material.

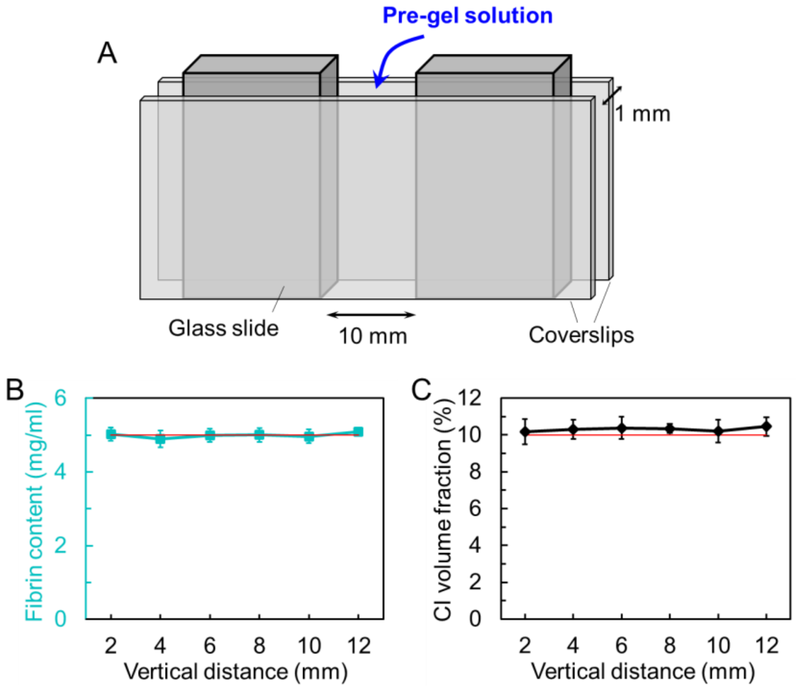


**Figure S18**. Preparation of fibrin gels embedded with CIPs. **A**. Custom chamber for vertical synthesis of fibrin gels with CIP inclusions. A custom-designed glass chamber was constructed using two 22x50 mm coverslips and two 25x15 mm glass slides as spacers. Coverslips and glass slides were treated with SurfaSil (Thermo Fisher) or 25% Triton X-100 to minimize the adhesion between the gel and the glass. The glass slides were placed between the coverslips, forming a 10 mm left-to-right gap to contain the pre-gel solution. The 1-mm thickness of each glass slide determined the fibrin gel thickness. The chamber was held together with binder clips (not shown) and positioned vertically on silicone grease to seal the bottom and prevent leaks. The pre-fibrin mixture was injected into the glass chamber after thrombin addition to initiate gelation. Once gelation was complete, the coverslips were removed. Due to CIP precipitation, the top portion of the gel (~2-3 mm) contained fewer CIPs, whereas the bottom portion (~1 mm) accumulated more CIPs, while the middle region was expected to have a uniform CIP distribution. Therefore, after gel formation, the top and bottom parts were discarded, and only the middle region was used for further experiments. This region was then cut into an 8 mm diameter sample for rheology measurements. **B** and **C**. Verification of uniform fibrin (B) and CIP (C) content across the vertically synthesized gel. The red line indicates the expected fibrin and CIP concentrations (5 mg/ml and 10% v/v, respectively). After full polymerization, the top and bottom ~4 mm sections were discarded, and the middle 12 mm section was cut into six 2-mm-wide strips, each corresponding to a 20 μl volume. Each strip was fully degraded with 0.25 mg/ml trypsin, and the fibrin concentration was determined via UV absorption, using degraded fibrinogen solutions as references. The CIP content was measured by washing the precipitate after degradation, drying, and weighing it. The density of CIPs in water was measured to be 6.92 g/ml, allowing determination of their corresponding volume fraction in the strips. The results confirm that both fibrin and CIPs are uniformly distributed inside the fibrin gel. The CIP content is slightly higher than the expected value, possibly due to oxidation during drying before weighing.

**Supplementary video 1**. Non-affine motions of coffee ground particles under compression, corresponding to Fig. 5D. Scale bar = 25 μm.

**Supplementary video 2**. Fiber stretching due to dislocation between two particle clusters, corresponding to Fig. S16A. Scale bar = 25 μm.

**Supplementary video 3**. Fiber stretching due to rotational motion of a single particle, corresponding to Fig. S16B. Scale bar = 25 μm.

**Supplementary video 4**. Fiber stretching due to collapse of a particle cluster concurrent with percolation, corresponding to Fig. S16C. Scale bar = 25 μm.
